# Investigation of Thyroid Hormone Associated Gene-Regulatory Networks during Hepatogenesis using an Induced Pluripotent Stem Cell based Model

**DOI:** 10.1101/2021.06.09.447730

**Authors:** Audrey Ncube, Nina Graffmann, Jan Greulich, Bo Scherer, Wasco Wruck, James Adjaye

## Abstract

Currently, the only treatment of end-stage liver diseases is liver transplantation. The worldwide shortage of donor organs and the high number of patients suffering from end-stage liver diseases, require alternative treatment approaches. Therefore, the generation of hepatocyte-like cells (HLCs) derived from induced pluripotent stem cells (iPSCs) is a promising treatment option. HLCs are suitable for patient-specific drug screening, disease modeling and regenerative medicine. So far, they are immature and resemble the fetal state. In this study, we employed the combined thyroid hormones triiodo-L-thyronine (T3) and L-thyroxine (T4) to drive HLCs towards maturation. HLCs expressed the maturation markers ALBUMIN (ALB), ALPHA-1 ANTITRYPSIN (A1AT), CYTOCHROME P450 3A4 (CYP3A4) and TRANSTHYRETIN (TTR). Remarkably, stimulation with T3 and T4 slightly reduced the expression of the fetal marker CYTOCHROME P450 3A7 (CYP3A7) and had an even more pronounced effect on ALPHA-FETOPROTEIN (AFP) levels. Comparative transcriptome and associated pathways in unstimulated and T3 and T4 stimulated HLCs revealed regulated expression of numerous genes within for example the peroxisome proliferator-activated receptor (PPAR), transforming growth factor beta (TGF-β), mitogen-activated protein kinase/extracellular-signal-regulated kinase (MAPK/ERK) signaling pathways and thyroid hormone synthesis. We propose the inclusion of combined T3 andT4 in HLC differentiation protocols to enhance maturation and therefore provide additional improvement in their applications in drug screening and disease modeling.

## 1. Introduction

The adult liver, being the largest internal organ of the human body, is organized in four lobes and consists mainly of hepatocytes [1]. Origin of the liver during development lies within the endoderm, one of the three germ layers. It is highly involved in metabolism by detoxification, protein synthesis, and glucose and fatty acid metabolism [2].

Liver diseases are a major threat, and research leading to treatment of these diseases is highly important. Although the liver is capable of regenerating itself, making it unique in the field of organ regeneration, the regeneration ability is also limited and compromised, especially when it comes to end-stage liver diseases. Hepatocytes re-enter the cell cycle and proliferate into new liver tissue, therefore not regenerating in a classical way, but compensating by building new liver mass [3]. The increasing number of liver diseases and the limited number of donor organs increase the importance of discovering optimal medical treatment plans for liver diseases. According to the regeneration potential of the liver, transplanting hepatocytes could also be used as treatment method, as the transplanted hepatocytes would replace the damaged liver tissue [4]. Immune rejection and insufficient isolation from donor liver tissue are problems occurring during treatment of liver diseases [5].

Primary human hepatocytes (PHH) are used as in vitro models of the liver, but difficulties arise in deriving and cultivating PHHs in vitro, as proliferation conditions are not yet completely understood [6]. To circumvent this problem, renewable sources for generating liver tissue have been established, resulting in the development of protocols for pluripotent stem cell-derived hepatocyte-like cells (HLCs) in two- and three dimensional cultures. Yamanaka et al. were able to generate induced pluripotent stem cells (iPSCs), thereby overcoming issues of immune rejection when patient specific cells are used or ethical issues faced with using embryonic stem cells [7]. This paved new paths in the field of regenerative medicine and disease modeling [8]. Many attempts to generate HLCs from ESCs and iPSCs using various small molecules and factors have been reported [9]–[12]; Varghese et al. compared growth factor- and small molecule-induced differentiation protocols to find the most robust and cost-effective protocol [10]. Of late, protocols involving 3D cultures are on the rise, because 2D cultures are limited by de-differentiation and lack interactions between cells and extracellular niche, thereby limiting clinical application [13], [14]. Although the liver is not rich in extracellular matrix (ECM), ECM is crucial in maintaining the differentiated state of hepatocytes by inducing intracellular signaling, hence it also plays a role in interactions within hepatocytes and maintaining functionality and polarity in 3D models [6]. Hepatic 3D cultures have been shown to resemble the liver more in terms of phenotype and gene expression [6]. Rashidi et al. showed the importance of differentiation in 3D, they generated 3D liver organoids that were kept in culture for over a year and still exhibiting CYP3A activity [13]. So far published protocols, both 2D and 3D, have not achieved significant down-regulation of the fetal markers-AFP and CYP3A7.

In this study, we address the issue of generating mature HLCs exhibiting the same functionality as liver biopsy-derived hepatocytes. We observed down-regulated expression of the fetal marker AFP and to a lesser extent also CYP3A7. Based on this, we propose the addition of synthetic analogs of the thyroid hormones triiodo-L-thyronine (T3) and L-thyroxine (T4) in the cell culture medium. Thyroid hormones are normally produced by the thyroid gland [15].

## 2. Materials and Methods

### 2.1. Culture of pluripotent stem cells

A previously established urine SIX2-progenitor cell derived induced pluripotent stem cell (iPSC) line ISRM-UM51 [16] and embryonic stem cell (ESC) line H9 (WiCell, Madison, WI, USA) were used for this experiment. The pluripotent stem cells (PSCs) were maintained on Matrigel^®^-coated dishes. For cultivation, StemMACS iPS-Brew XF Medium with 1% Penicillin/Streptomycin (P/S) was used. The medium was changed on a daily basis and at a confluency of about 80%; the cells were harvested and split. For this, the iPS cells were incubated with PBS (-/-) for seven minutes at room temperature and then scrapped off the plate and the cell suspension was centrifuged at 40 G for five minutes.

#### 2.1.1. Ethics statement

The use of iPSC lines for this study was approved by the ethics committee of the medical faculty of Heinrich-Heine University under the study number 5704.

### 2.2. Differentiation of PSCs to Hepatocytes

Dose dependency of the thyroid hormones was confirmed by preliminary experiments with HepG2 cells according to Conti et. al, who showed that addition of the thyroid hormones at a concentration of 10^−6^ or 10^−8^ M suppressed AFP secretion. The iPS and ES-H9 cells were differentiated to hepatocyte-like cells (HLC) following our previously described and slightly modified three step protocol [12], [17], [18]. In the first step, the cells were differentiated into the definitive endoderm (DE), using DE medium (96% RPMI 1640, 2% B27 without retinoic acid, 1% GlutaMAX^®^, 1% P/S and supplemented with 100 ng/mL Activin A (Peprotech, Hamburg, Germany, 1:1000) and CHIR99021 (Tocris, Wiesbaden-Nordenstadt, Germany, 20 mM) (added only on the first day, 1:8000)). The second step drives the cells into hepatic endoderm (HE), cells are cultured from day 4 to 7 in HE medium consisting of 78% Knock-Out DMEM, 20% Knock-Out Serum, 0.5% GlutaMAX^®^, 1% P/S, 0,01% 2-Mercaptoethanol (all Gibco) and 1% DMSO (Sigma). Finally, for the maturation step (82% Leibovitz15 Medium, 8% fetal bovine serum, 8% Tryptose Phosphate Broth, 1% P/S, 1% GlutaMAX^®^ (all Gibco) and 0.06% Insulin (Sigma), supplemented with hepatocyte growth factor (HGF) (Peprotech 10 ng/mL), dexamethasone (DEX) (25 ng/mL) were used. For the first two steps medium was changed every day, and then every second day in the last step. T3/T4 were included in the maturation medium from day 8 onwards. One milliliter cell culture supernatant was collected from each stage DE, HE and HLC different conditions before medium change for UREA or ELISA assays and stored at −20°C.

In a first differentiation, UM51-derived HLCs were incubated with either 10^−6^ M T3 or a combination of 10^−6^ M T3 and T4 respectively. In a second differentiation, to compare the effects of T3 and T4 on the differentiation of iPSCs and ESCs, a combination of 10^−6^ M T3 and T4 was added to the HLC medium of both UM51- and H9-derived HLCs during the maturation stage.

#### 2.2.1. Immunocytochemistry

For immunofluorescence staining, cells were fixed with 4% Paraformaldehyde for fifteen minutes, followed by two hours blocking of unspecific binding sites using blocking buffer composed of 10% normal goat serum (NGS, Sigma), 1% bovine serum albumin (BSA, Sigma), 0.5% Triton (Carl Roth GmbH & Co. KG, Karlsruhe, Germany) and 0.05% Tween 20 (sigma), all dissolved in PBS. If the target stain was extracellular, Triton and Tween were excluded from the blocking buffer. Primary antibodies were incubated overnight at 4°C; AFP (1:200), ALB (1:500) (Sigma-Aldrich Chemie), HNF4α (1:250; Abcam), SOX17 (1:50; R&D Systems), E-CAD (1.200; Cell Signaling Technology^®^) followed by one hour secondary antibody incubation; Alexa488 and Cy3 (Thermo Fischer Scientific, 1:2000), as well as Hoechst 33258 dye (Sigma-Aldrich, 1:5000) for cell nuclei staining. Imaging was done using a fluorescence microscope (LSM700; Zeiss, Oberkochen, Germany) and the ZenBlue 2012 Software Version 1.1.2.0 (Carl Zeiss Microscopy GmbH, Jena, Germany) for picture processing.

#### 2.2.2. RNA Isolation and cDNA synthesis

The Direct-zol RNA Miniprep Kit (Zymo Research, CA, USA) was used for RNA isolation according to manufacturer’s instruction. Five hundred nanogram of the isolated RNA were reverse transcribed into complementary DNA (cDNA) synthesis using the TaqMan Reverse Transcription (RT) Kit (Applied Biosystems). The reaction mixture (10μl per sample) included 3.85 μL H2O, 1 μL reverse transcriptase buffer, 2.2 μL MgCl2 (25 mM), 0.5 μL Oligo (dT)/ Random hexamer (50 μM), 2μL dNTP mix (10 mM), 0.2 μL RNase Inhibitor (20 U/μL) and 0.25 μL Reverse Transcriptase (50 U/μL).

#### 2.2.3. Quantitative real-time polymerase chain reaction (qRT-PCR)

Quantitative real-time PCR was carried out in a in a VIIA7 machine (Life technologies), each sample in triplicate. One μL cDNA of each sample was placed in each well and 9 μL of the Master Mix consisting of Power SYBR Green Master Mix (Life technologies), water and primers purchased from MWG. The specific primer sequences are provided in supplementary table S2. The housekeeping gene ribosomal protein S16 (RPS16) was used for normalization of the tested genes. The program started with denaturation of the samples at 95°C for two minutes, followed by forty cycles amplification with a thirty seconds denaturation step at 95°C, annealing step at the primer-specific temperature (57°C to 63°C) for thirty seconds and extension at 72°C for thirty seconds. The expression levels were calculated using the ΔΔCT method and are shown as mean values with standard error of mean.

#### 2.2.4. Western blot analysis

For the protein extraction, cells were harvested and lysed in RIPA buffer (Sigma Aldrich) supplemented with a complete protease and phosphatase inhibitors cocktail (Roche, Sigma), and protein quantified using the Pierce BCA Kit. Further, cell lysates were loaded on a 4 - 20% Bis-Tris gel (Invitrogen) and proteins blotted onto a 0.45 μm nitrocellulose membrane (GE Healthcare Life Sciences). The membranes were then blocked with 5% skimmed milk in Tris-Buffered Saline Tween (TBS-T) and incubated overnight at 4°C with the respective primary antibodies from Sigma-Aldrich Chemie: ALB (1:2500 mouse, TBS-T 5% Milk) and AFP (1:2000 rabbit, TBS-T 5% Milk) or from CST: β-actin (1:5000 mouse, TBS-T 5% Milk). For the detection HRP (horseradish peroxidase) coupled secondary antibodies ((1:4000) were used and detected via chemiluminescence on a Fusion FX instrument (PeqLab). Signals were quantified with Fiji (Image J) and normalized to β-actin. Significance was calculated by ANOVA.

#### 2.2.5. Fluorescence activated cell sorting (FACS)

For flow cytometric analyses, cells were harvested with trypsin, fixed for 15 min with 2% PFA and permeabilized with 0.5% Triton in PBS. Primary antibody incubation was carried out overnight. After 3 times washing, cells were incubated for 1h in the dark. Flow cytometric analysis was performed on a CyAn and analyzed with the summit software.

### 2.3. Hepatocyte functionality tests

#### 2.3.1. Indocyanine Green (ICG) clearance test

Here we aimed at investigating the uptake and release of indocyanine green (ICG) by the HLCs as a measure of active transport. The dye, Cardiogreen (1 mg/mL) in HLC medium was added in one well and the cells incubated for thirty minutes at 37°C. Afterwards, cells were washed with PBS (+/+) and medium changed. ICG clearance was observed over a period of twenty hours and images captured under a light microscope.

#### 2.3.2. Urea Assay

The QuantiChrom™ Urea Assay Kit was used to measure Urea secretion of the HLCs. Media from the stages DE, HE and HLC were taken from all conditions of the cells and stored at −20°C. The urea assay was carried out according to the manufacturer’s instructions. Final concentrations of secreted urea were calculated and analyzed using the prepared standard curve.

#### 2.3.3. Glycogen storage

First, cells were fixed with 4% PFA and then treated with periodic acid solution for 5 minutes, followed by several washing steps with water. Afterwards, cells were incubated in Schiff’s Reagent for fifteen minutes and then the cell nuclei stained with hematoxylin for ninety seconds. Finally, cells were washed again with water and images captured under a light microscope.

### 2.4. Transcriptome analysis

HLCs treated with T3/T4 hormones and untreated as well as commercial fetal and adult liver samples were submitted to the BMFZ (Biomedizinisches Forschungszentrum) core facility of the Heinrich-Heine Universität, Düsseldorf for microarray hybridization on the Affymetrix Human Clariom S assay. Raw data (CEL files) were read into the R/Bioconductor environment [19]. Background-correction and normalization via the Robust Multi-array Average (RMA) method were achieved using the R package oligo [20]. The dendrogram was calculated via the R hierarchical clustering function hclust using Pearson correlation and complete linkage agglomeration. Gene expression was determined with a detection-p-value threshold of 0.05 while the detection-p-value was assessed according to the method from Graffmann et al. [18]. Venn diagrams were drawn based on the gene expression determined this way with the R VennDiagram package [21].

#### 2.4.1. Analysis of pathways and Gene ontologies (GOs)

KEGG (Kyoto Encyclopedia of Genes and Genomes) [22] pathways and associated genes were downloaded from the KEGG website in March 2018. Overrepresented pathways were determined via the R-built-in hypergeometric test. Up- and down-regulated genes in the KEGG PPAR signaling pathway chart were marked in red (up-regulation) and green (down-regulation) via the R pathView package [23] using log2-ratios between T3/T4-treated and untreated HLCs. Overrepresented GOs were detected employing the R package GOstats [24].

#### 2.4.2. Data Availability

All microarray data will be made available at NCBI GEO under the accession number XXX when the manuscript is accepted.

## 3. Results

### 3.1. HLC Differentiation

The iPSC- line UM51 was differentiated into HLCs following our modified three step protocol [12], [17], [18]. HepG2 cells were treated for seven days and medium was changed on every second day. Afterwards, immunofluorescence stainings for ALB and AFP were performed, as well as a qRT-pCR and a Western blot. From these results we concluded that 10^−6^ M is the best concentration for the treatment-figure S1. At day eight of the differentiation protocol, T3 and a combination of T3 and T4 were added to the differentiation medium. In a second experiment, differentiation efficiency between iPSCs and ESCs was compared. The iPSC- line UM51 and the ESC-line H9 were differentiated into HLCs in the same way, but comparing standard HLC condition versus addition of the combination of T3 and T4. The differentiations were stopped after eighteen days. Phase contrast images of the cells were captured at distinct stages throughout the differentiation in order to analyze changes in cell morphology, Figure 1. At the pluripotent stage (day 0), the cells showed the typical sharp edged and well defined colonies. During the differentiation, typical cell morphology changes were observed in the DE, HE and HLC stages, resulting in cobblestone epithelial shaped cells. In addition to the phase contrast images, cells of all stages were stained for the expression of typical markers of each stage (Fig 1). In the DE stage, the cells expressed the transcription factor SOX17, which is involved in cell fate decision and plays a key role in embryonic development. In the HE stage, most cells expressed the transcription factor HNF4α, which is essential for liver development, and some cells expressed AFP, which is a fetal marker of the liver, usually expressed only during development.

**Figure 1.**
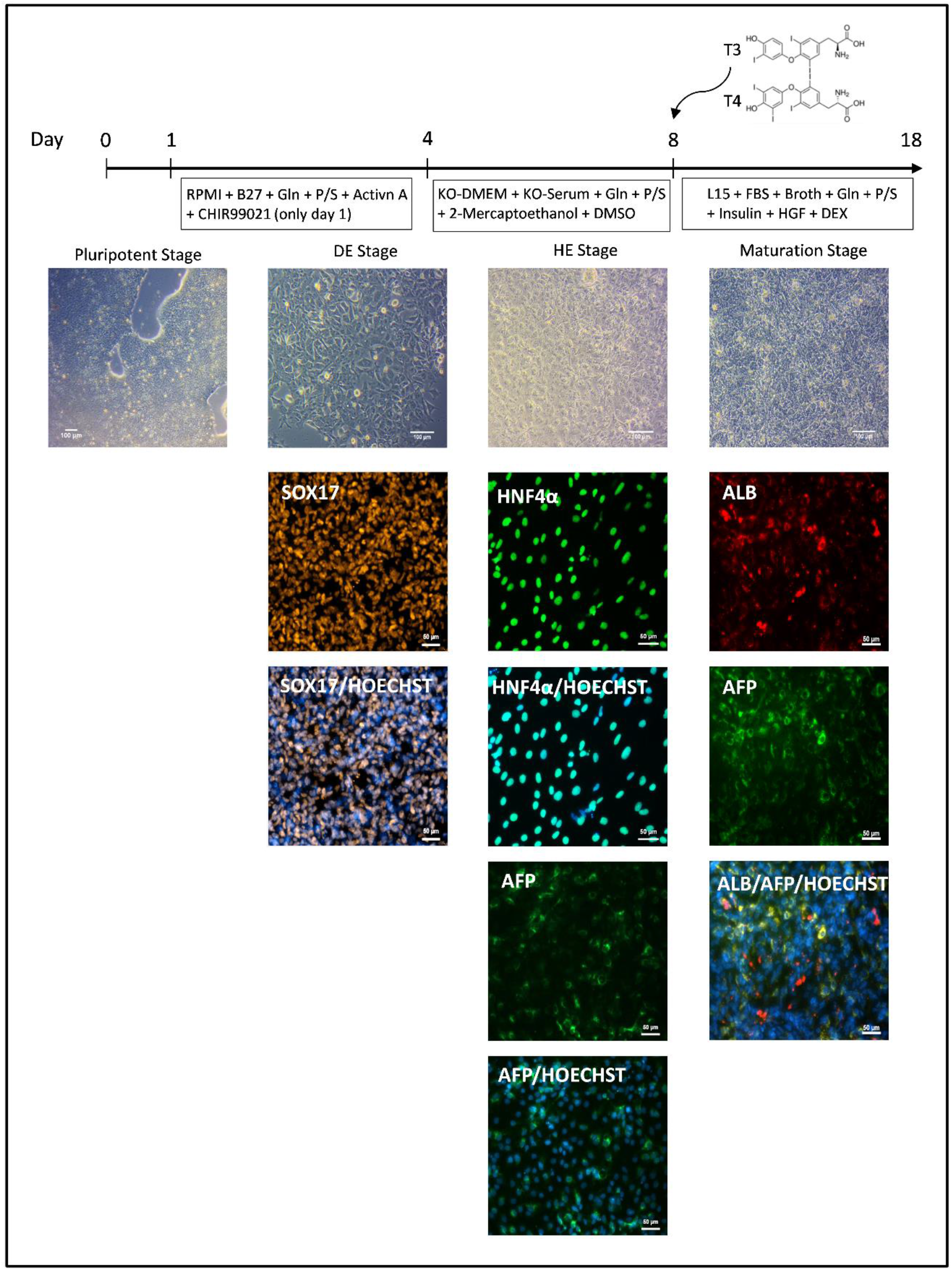
Differentiation scheme of pluripotent stem cells into hepatocyte-like cells. For each stage of differentiation, a phase contrast picture of the cells and a staining specific for each stage is provided. Beginning at day eight, T3 and T4 were included in the differentiation medium until the end. (Bright-field images Scale bars = 100 μm, immunostaining images scale bars = 50 μm).

At the maturation stage, after eighteen days of differentiation, up to 85% of HLCs expressed ALB (Fig, 1, S2), which is known as a maturation marker in the liver, however, a few cells still expressed the fetal marker AFP (Fig. 1). ALB and AFP immunostaining images comparing HLCs generated under standard condition and those treated with T3 and T4 are shown in supplementary Figure S3.

#### 3.1.1. HLC functionality assays

In order to confirm the generation of HLCs, functionality assays were performed. The generated iPSC-derived HLCs cells were able to store glycogen, as seen by magenta color retained by the cells, Figure 2A. Furthermore, the cells were able to take up and release the green dye ICG, Figure 2B. The ICG clearance is a common test for liver functionality, as only mature and functional livers are able to take up and release ICG.

**Figure 2.**
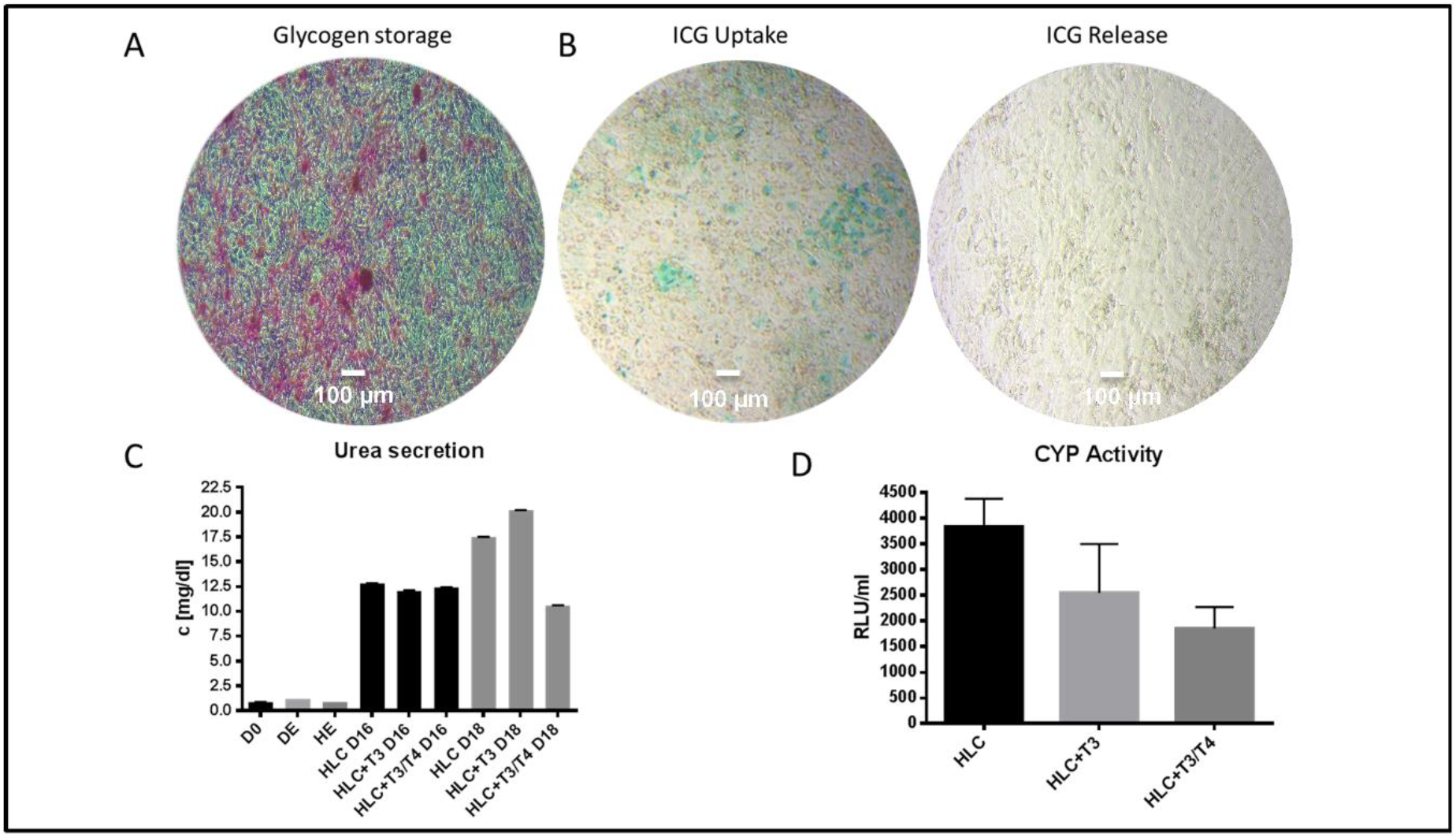
Functionality assays carried out for the generated HLCs. A glycogen storage assay, ICG Uptake-Release assay, urea assay and CYP-Activity assay for the cytochrome P450 3A4 were performed. (**A**) The cells were able to store glycogen. (**B**) The cells were able to take up ICG and release it completely after 18 hours. (**C**) Cells from day 0, DE stage and HE stage secreted nearly no urea, whereas HLCs from day 16 and day 18 of differentiation secreted all more than 10 mg/dl urea. (**D**) All generated HLCs showed activity of the CYP3A4. (Bright-field images Scale bars = 100 μm).

In addition, conditioned medium was collected from the cells and urea secretion measured. The HLCs in all conditions showed a urea secretion of more than 10 mg/dl after sixteen and eighteen days (Fig. 2C). Interestingly, longer differentiation seemed to decrease urea secretion in the HLC + T3/T4 condition after eighteen days. Moreover, after eighteen days of differentiation all HLCs showed cytochrome P450 3A4 activity, with the lowest activity observed in the HLC + T3/T4 condition (Fig. 2D).

#### 3.1.2. Confirmation of expression of T3 and T4 regulated genes

Gene expression of fetal and adult genes of the HLCs was analyzed by qRT-PCR. The fetal gene AFP was downregulated in iPSC-derived HLCs after addition of T3 and T3/T4 to the differentiation protocol, but only significantly in the T3 condition (Fig. 3A, B and C). Furthermore, the maturation markers ALB, CYP3A4, A1AT and TTR were expressed in all conditions. The fetal marker CYP3A7 was slightly downregulated in HLCs after addition of T3 and T3/T4 to the differentiation medium (Fig. 3A). In the parallel differentiation comparing UM51 and H9 derived HLCs, the fetal markers AFP and CYP3A7 were downregulated after addition of T3/T4, whereas ALB is expressed in both HLC preparations (Fig. 3B and C, Figure S3).

**Figure 3.**
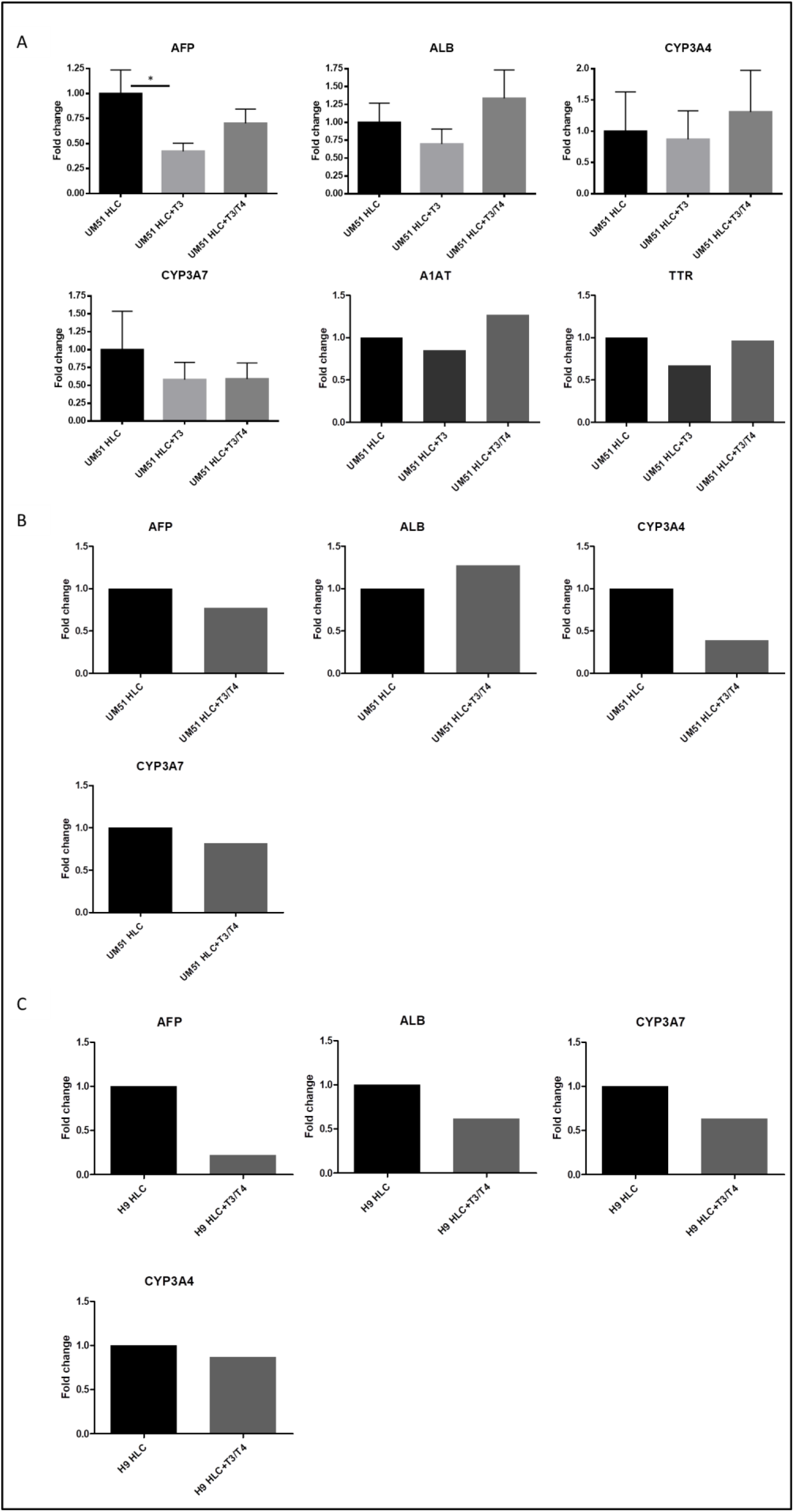
qRT-PCR carried out for the generated HLCs. (**A**) qRT-PCR for HLCs generated from iPSC-line UM-51 with primers used for AFP, ALB, CYP3A4, CYP3A7, A1AT and TTR. AFP was significantly downregulated after treatment with T3. n=3 for AFP, ALB, CYP3A4 and CYP3A7, n=1 for A1AT and TTR. (**B**) Second qRT-PCR for HLCs generated from iPSC-line UM-51 with primers used for AFP, ALB, CYP3A4 and CYP3A7 only treated with T3/T4. n=1. (**C**) qRT-PCR for HLCs generated from ESC-line H9 with primers used for AFP, ALB, CYP3A4 and CYP3A7, n=1. Error bars represent S.E.M., ^*^ = p < 0.05

### 3.2. Down regulated protein expression of AFP

In the iPSC-derived HLCs, the fetal marker AFP was significantly downregulated on the protein level after addition of T3 and T3/T4 to the differentiation protocol (Fig. 4A). In additional differentiation comparing ESC-derived and iPSC-derived HLCs by adding T3/T4, Western blot analysis revealed a downregulation of AFP both in ESC- and iPSC-derived HLCs after addition of T3/T4. In accordance with this observation, secreted AFP levels were also reduced after T3/T4 treatment of ESC derived HLCs (Supplementary Fig. S4).

**Figure 4.**
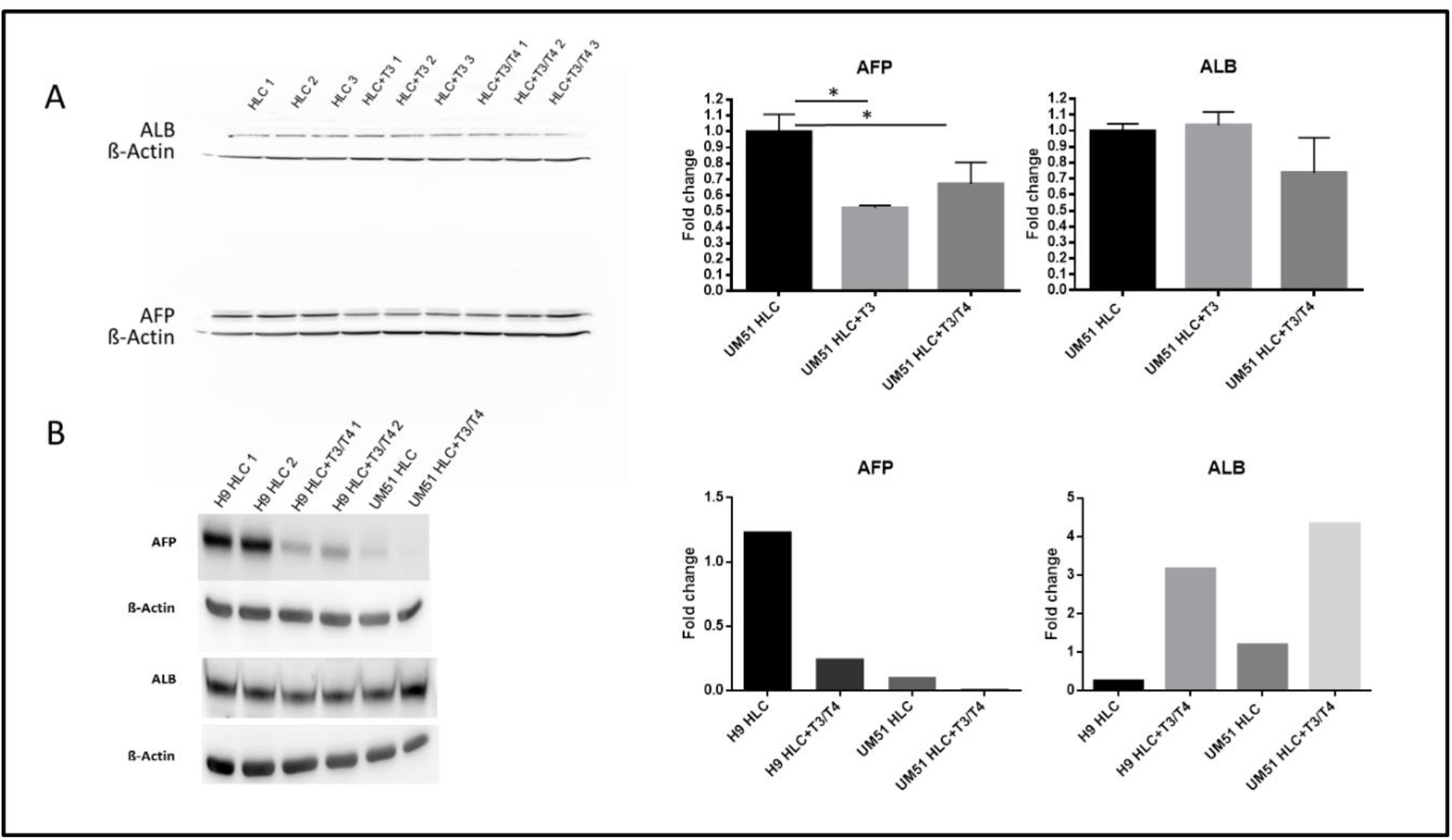
Western blot was performed for ALB and AFP for the HLCs. (**A**) Western blot for HLCs generated from iPSC-line UM-51. AFP was significantly downregulated after adding T3 and T3/T4 to the differentiation protocol. ALB showed no significant changes within the different conditions. n=3 (**B**) Western blot of HLCs generated from ESC-line H9 and iPSC-line UM-51. AFP was downregulated in both cell lines after addition of T3/T4, while ALB was upregulated. For H9 HLC and H9 HLC+T3/T4 n=2, for UM51 HLC and UM51 HLC+T3/T4- n=1, for evaluation n=1 for all samples for a better comparison. Error bars represent S.E.M., * = p < 0.05

### 3.3. Transcriptome analysis

ESC- and iPSC-derived HLCs with and without addition of T3/T4 were subjected to transcriptome analysis together with samples of fetal and adult liver as control. The dendrogram in Figure 5A resulting from global transcriptome cluster analysis with a coefficient of variation (cv) > 0.1 shows a cluster of adult and fetal liver and a cluster of HLCs with two sub-clusters determined by the original cell source. In Figure 5B, a table of Pearson correlation coefficients of gene expression data of all samples versus each other is depicted.

**Figure 5.**
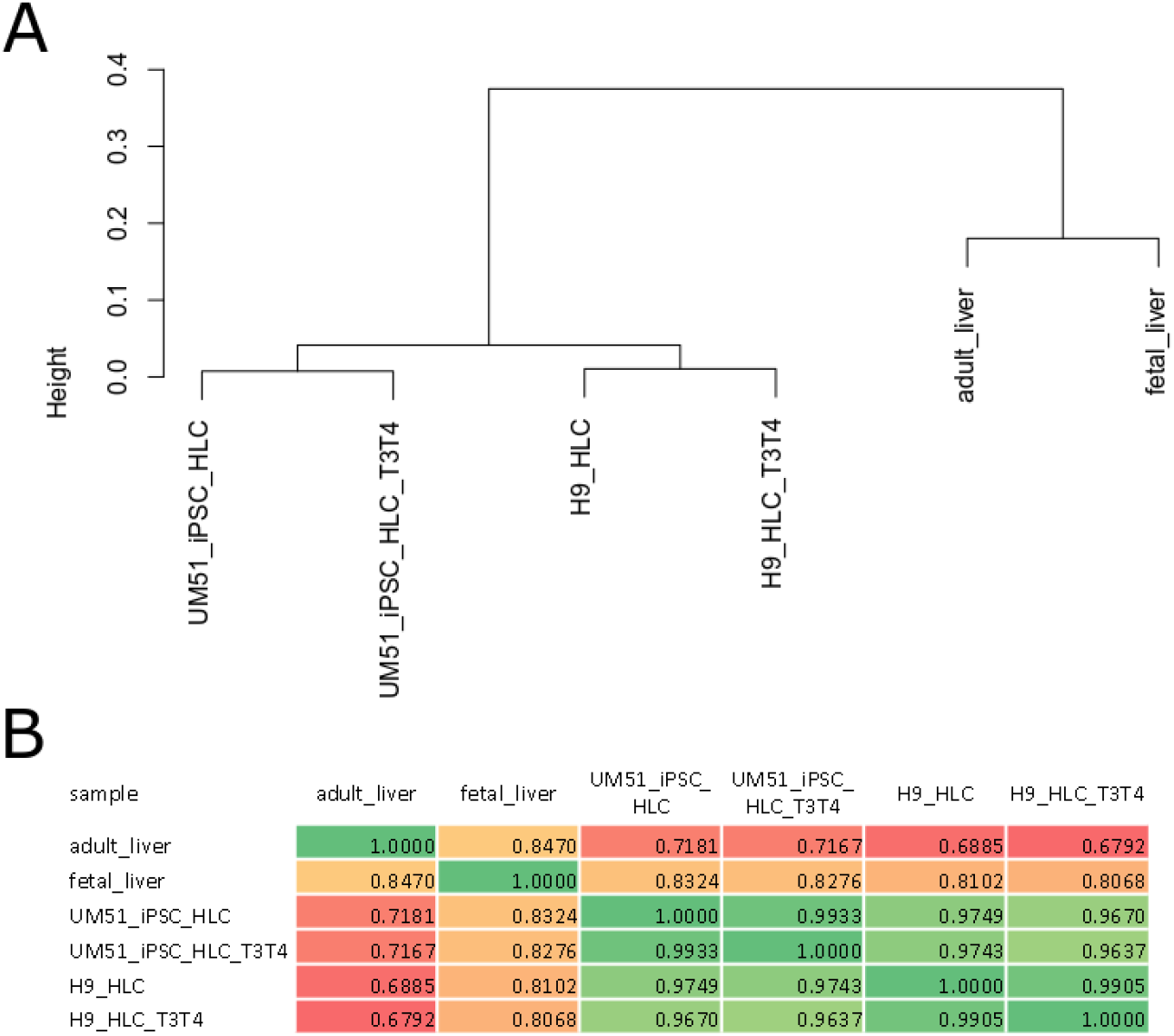
Global transcriptome cluster analysis and Pearson correlations. (**A**) The dendrogram results from global transcriptome cluster analysis of probesets with a coefficient of variation (cv) > 0.1 using Pearson correlation as similarity measure and complete linkage as cluster agglomeration method. (**B**) The table shows the Pearson correlation coefficients of gene expression data of all samples vs. each other.

Supplementary table S1 shows that the treatment of H9-HLCs with T3/T4 resulted in down-regulation of numerous metabolic pathways such as Drug metabolism - cytochrome P450 (p=0.0002, q=0.0031), Metabolism of xenobiotics by cytochrome P450 (p=0.0003, q=0.0031), Glutathione metabolism (p=0.0027, q=0.0108), Pyruvate metabolism (q=0.0437) and Glycolysis (p=0.0460, q=0.092). In treated UM51-iPSC-HLCs, down-regulated pathways were over-represented with a considerably higher significance due to a higher number of involved genes as shown in Supplementary table S1. Cholesterol metabolism was most significant (p=1.46E-07, q=4.2E-06), followed by Fat digestion and absorption (p=4E-05, q=0.0004), Complement and coagulation cascades (p=4.3E-05, q=0.0004) and vitamin digestion and absorption (p=0.0003, q=0.0018). In the pathways over-represented in the up-regulated genes in T3/T4-treated H9-HLCs compared to untreated H9-HLCs (Supplementary Table S1) we identified Thyroid hormone synthesis (p=0.032, q=0.0668). Furthermore, we found thyroid hormone generation (p=0.0009) in the GOs over-represented in up-regulated genes in T3/T4-treated UM51-HLCs compared to untreated UM51-iPSC-HLCs (Supplementary Table S1). We analyzed cytotoxicity at the transcriptome level by assessing genes from the KEGG pathway “Natural killer cell mediated cytotoxicity”. The heatmap in Supplementary Figure S5 demonstrates that there is no specific change in cytotoxicity-related gene expression induced by the T3/T4 treatment. Furthermore, we examined genes from the proliferation-related KEGG pathway “Cell cycle” at the transcriptome level and found that there were non-significant changes as shown in the heatmap Supplementary Figure S6.

Finally, we checked for GOs overrepresented in genes commonly up- or down-regulated in HLCs from H9 and UM51 cells after T3/T4 treatment (Supplementary Table S1). Interestingly, we found that mesenchymal cell differentiation was among the common down-regulated GOs. This might imply a higher stability within the epithelial HLC population as aberrant de-differentiation into a mesenchymal phenotype seems to be suppressed by T3/T4.

#### 3.3.1. Analysis of the PPAR signaling pathway

We investigated in more detail the peroxisome proliferator-activated receptor (PPAR) signaling pathway in which we found down-regulated genes in the T3/T4-treated H9-HLCs and UM51-iPSC-HLCs (Supplementary Table 1). PPAR signaling plays a key role in the liver [25] - not only by regulating metabolism, but also by modulating cell differentiation, proliferation, apoptosis, and aging. The addition of T3/T4 to HLCs in our experiments down-regulates most genes in the PPAR signaling pathway. The KEGG pathway chart in Figure 6 shows up- or down-regulation of several genes in untreated UM51-iPSC-derived HLCs (left part of the gene boxes) and T3/T4-treated UM51-iPSC-derived HLCs (right part of the gene boxes). Reciprocal regulation between Thyroid receptors (TR) and PPARs has already been reported for rodents by Flores-Morales et al. [26] and Lu and Cheng [27].

**Figure 6.**
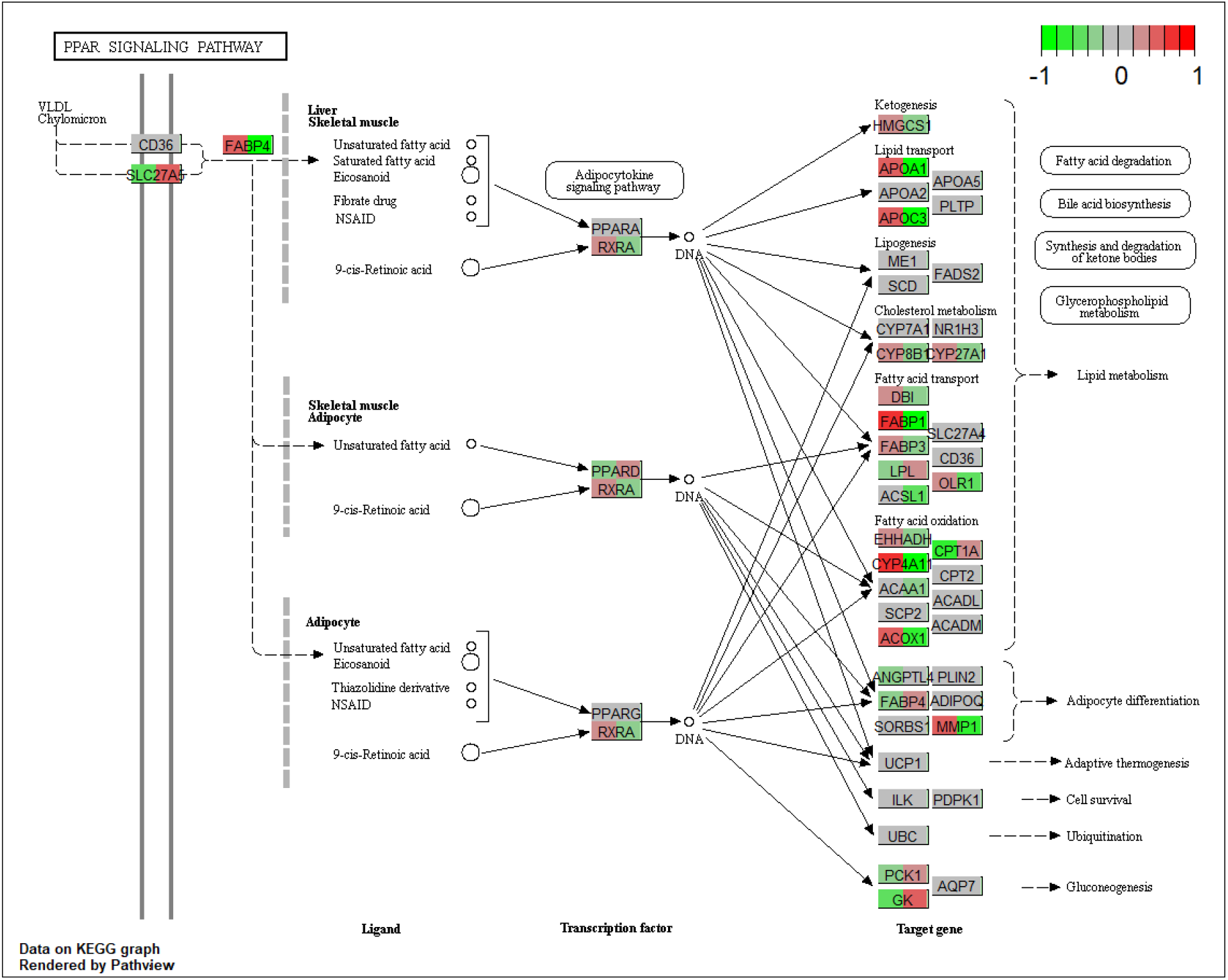
Pathway chart of PPAR signaling pathway indicating T3/T4-mediated up- or down-regulation. The chart shows up- or down-regulation of genes in untreated UM51-iPSC-derived HLCs (left part of the gene boxes) and T3/T4-treated UM51-iPSC-derived HLCs (right part of the gene boxes). Log2-ratios were calculated by taking log2 of the quotient of the expression of HLC or T3/T4-treated HLC divided by the mean of HLC and T3/T4-treated HLC. The color scale is located in the top right of the chart, red indicates up-regulation, green down-regulation.

#### 3.3.2. Comparison of T3/T4-treated HLCs with adult and fetal liver

We set out to further elucidate the gene expression patterns of the distinct conditions tested in our experiments in more detail. With this intention, we dissected expressed genes filtered via a threshold for a detection-p-value < 0.05 in a Venn diagram comparing the four conditions of fetal liver, adult liver and HLCs untreated and treated with T3/T4 thyroid hormones (Figure 7A). Of most relevance we considered genes which were expressed in common in T3/T4 treated HLCs and adult liver. Our hypothesis was that by the overlap with the adult phenotype these genes would indicate maturation of the HLCs induced by the T3/T4 thyroid hormone treatment. Nineteen genes were expressed in T3/T4-treated HLCs and in adult liver in common (marked in yellow). These nineteen genes were subjected to a KEGG pathway over-representation analysis (Figure 7B) which revealed inflammatory mediator regulation of TRP channels as most significant pathway. There is only limited knowledge on the role of TRP channels in liver including roles in cell proliferation, migration, lysosomal Ca(2+) release and development and progression of liver cancer [28]. The cold receptor TRPM8 which we identified here has been reported to be expressed in liver and mutations in TRPM8 have been associated with impaired lipid levels [29] which are obviously connected to genuine liver functionality. Figure 7C lists GOs over-represented in the nineteen genes expressed in common in in T3/T4-treated HLCs and in adult. Here, the results include two GOs related to MAPK/ERK signaling and the GO developmental maturation. MAPK signaling has been implicated in liver maturation particularly via p38α MAPK [30], [31]. Looking into the gene lists of these GOs most outstanding is the gene NODAL – a part of the TGFβ-signaling pathway - which was described to force cell fate into mesoderm lineage upon low NODAL expression and into endoderm upon high expression [9], [32]. An auto-regulatory loop reinforces NODAL expression and endoderm specification [32].

**Figure 7.**
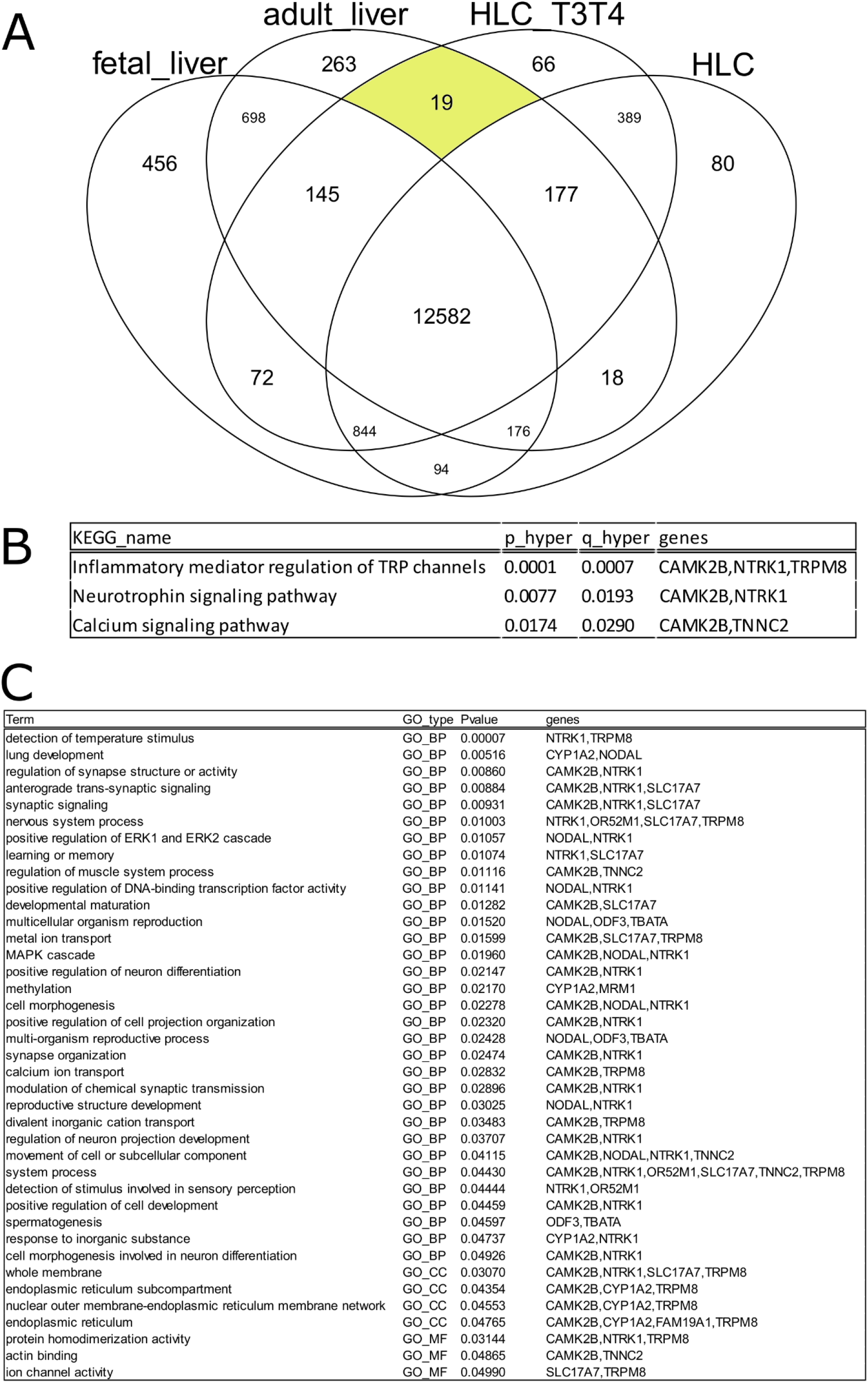
Venn diagram comparison of gene expression in untreated and T3/T4-treated HLCs with adult and fetal liver. (**A**) Genes expressed (detection-p-value < 0.05) in untreated HLCs, T3/T4-treated HLCs, adult liver and fetal liver are compared in a venn diagram. Nineteen genes are expressed in T3/T4-treated HLCs and in adult liver in common. (**B**) KEGG pathways over-represented in the nineteen genes expressed in common in in T3/T4-treated HLCs and in adult. (**C**) Gene ontologies over-represented in the nineteen genes expressed in common in in T3/T4-treated HLCs and in adult.

## 4. Discussion

Liver diseases are a major threat and a common cause of mortality [33]. To date, the only effective treatment for end-stage liver diseases is liver transplantation, but this is compromised by the low availability of donor organs, as well as immune rejection after transplantation. This has led to increased demand to find new alternative treatment options [5], especially in the field of regenerative medicine. The establishment of iPSC-derived hepatocyte-like cells has presented an enormous opportunity, as these cells can be used in patient-specific disease modeling and drug screening and development [34], thereby circumventing the risk of immune rejection after transplantation [4], [8]. The only problem arising from this approach is the ability to generate mature and functional iPSC-derived hepatocyte-like cells which parallel liver-biopsy derived primary hepatocytes [34].

In this study, we focused on this aspect of hepatocyte-like cell maturation and investigated if this could be enhanced by stimulating the cell cultures with thyroid hormones T3 and T4. T3 is the mainly active and more potent form of thyroid hormones and derived from de-iodination of T4. T4 is the major occurring form of the thyroid hormone in the blood and referred to as a storage form of T3. The conversion of T4 to T3 takes place directly in the target tissues. The functions of T3 and T4 are mainly regulation of metabolism, as well as they play an essential role in development, differentiation and maturation of nearly all cells of the human body[15], [35]. The liver is known to be one of the main target organs for the thyroid hormones; the high interdependency between the thyroid gland and the liver results in risk of liver diseases when thyroid hormone levels are altered in an organism [36]. Two well described target genes of thyroid hormones are ALB, a liver maturity marker, and AFP, a fetal liver marker. Thyroid hormones have been shown to upregulate ALB and downregulate AFP in HepG2 cells [37], [38]. Tarim et al. showed that T3 promotes hepatocyte proliferation mediating organ growth, whereas in other organs T3 induced differentiation of several progenitor cells, thereby mediating cell maturation in the target organ [39]. As already mentioned above, the conversion of T4 to T3 takes place directly in the target tissues. Cvoro et al. also showed that T3 induced KLF9 in HepG2 cells. They also treated ESCs and iPSCs differentiated into definitive endoderm and terminally differentiated iPSC-derived hepatocytes with T3 and only showed successful KLF9 induction [40].

By adding T3 and T4 to the differentiation medium of pluripotent stem cell-derived HLCs, we aimed to enhance maturation of the generated HLCs. Our main focus was on the upregulation of the maturity marker ALB and the downregulation of the fetal marker AFP. After differentiation, cells were successfully stained for various liver markers; SOX17 at the DE stage, HNF4α and AFP at the HE stage and ALB and AFP at the HLC stage (Fig. 1). The hepatocyte functionality tests-glycogen storage, ICG clearance, urea secretion and cytochrome activity assay-also confirmed maturation to a certain degree of the generated HLCs (Fig. 2). Of outmost interest for us was the mRNA and protein level expression of ALB and AFP. Indeed, our results show down-regulation of AFP expression after addition of the thyroid hormones compared to the standard HLC protocol without thyroid hormones. The down-regulation could be seen on both mRNA (Fig. 3) and protein levels (Fig. 4), thus indicating a stable decrease of AFP mediated by the thyroid hormones. On the mRNA level, we could also observe the expression of A1AT and TTR in the UM51-derived HLCs (Fig. 3A), which are two maturity markers of the liver. Although there was no significant change in gene expression after adding thyroid hormones to the differentiation, expression of these two genes in all iPSC-derived HLCs indicates successful differentiation into HLCs and maturation to a certain extent. Although we could not observe a significant difference in control HLCs and those differentiated with T3 or T3/T4, our data suggests that the addition of T3 to the differentiation slightly increases maturity and functionality in pluripotent stem cell-derived HLCs in terms of cytochrome activity (Fig. 2D) and urea secretion, especially after eighteen days of differentiation (Fig. 2C), indicating that a longer differentiation may also increase maturity of HLCs.

Interestingly, when comparing iPSC- and ESC-derived HLCs, our data strongly suggests that urine-derived iPSCs seem to be more suitable for generation of mature HLCs. Western blot analysis revealed lower levels of the fetal marker AFP and higher levels of the maturity marker ALB in urine-derived iPSC-derived HLCs compared to ESC-derived HLCs. AFP could be further downregulated by addition of thyroid hormones (Fig. 4 B).

Detailed transcriptome analysis revealed that in line with reports mainly based on rodents T3/T4 treatment has a reciprocal effect on PPAR signaling [25]. Further dissecting the overlap between the T3/T4 treated cells and the mature phenotype indicated at MAPK-signaling and also the NODAL branch of TGFβ-signaling as potential actors driving maturation. These findings were underpinned by previous studies reporting possible roles of MAPK in liver maturation [30], [31] and the well-settled role of NODAL in early liver development [32], but need further evaluation in larger studies. In addition, we saw that genes commonly reduced after T3/T4 treatment in H9 and UM51 derived HLCs were associated with mesenchymal cell development. Dedifferentiation of in vitro derived HLC into the mesenchymal lineage is a well-known problem of this process. Sometimes activation of the TGFβ pathway with small molecules is employed to counteract this phenomenon which might be dispensable in the presence of T3/T4.

## 5. Conclusions

Our findings have revealed that the thyroid hormones T3 and T4 downregulate the expression of the fetal liver markers and AFP and to a lesser extent CYP3A7 thereby driving pluripotent stem cell-derived hepatocyte-like cells a step closer towards maturation.

## Supplementary Materials

**Table S1.** Gene list, GO analysis and KEGG pathways comparing H9 vs UM51 HLCs and T3T4 treated vs untreated HLCs. **Table S2**. List of specific primer sequences used for the quantitative real-time PCR. **Figure S1**. Concentration determination of T3 and T4 after 7 days treatment of HepG2 cells. **Figure S2:**Representative flow cytometric analysis of ALB expression in HLCs. Blue curve: secondary antibody control, green curve: ALB staining **Figure S3**. Immunostaining images of HLCs generated under standard condition compared to those generated under T3/T4 treatment, showing ALB and AFP staining. **Figure S4:**Western blot was performed with concentrated supernatants from ESC derived HLCs. (**A**) AFP could be detected in the supernatant. (**B**) Ponceau S staining. **Figure S5**: Genes from the KEGG pathway “Natural killer cell mediated cytotoxicity” (hsaO465O) are not affected by the T3/T4 treatment. **Figure S6**: Genes from the KEGG pathway “Cell cycle” (hsa04110) are only marginally affected by the T3/T4 treatment.

## Author Contributions

Conceptualization: J.A. and A.N.; Methodology, Investigation, Validation: A.N., J.G. and B.S.; Formal analysis N.G., A.N., J.A. and W.W.; Data curation: N.G and W.W.; Writing-original draft preparation: A.N., J.G. and W.W.; Writing – review and editing: J.A., W.W., N.G. and A.N.; Supervision: J.A.

## Acknowledgments

James Adjaye acknowledges support from the medical faculty of Heinrich Heine University-Düsseldorf, Germany.

## References

1. K. Vdoviakováet al., “Importance Rat Liver Morphology and Vasculature in Surgical Research.,” Med. Sci. Monit, vol. 22, pp. 4716–4728, Dec. 2016, doi: 10.12659/MSM.899129.

2. D. P. Bogdanos, B. Gao, and M. E. Gershwin, “Liver immunology.,” Compr. Physiol., vol. 3, no. 2, pp. 567–98, Apr. 2013, doi: 10.1002/cphy.c120011.

3. S. A. Mao, J. M. Glorioso, and S. L. Nyberg, “Liver regeneration.,” Transl. Res., vol. 163, no. 4, pp. 352–62, Apr. 2014, doi: 10.1016/j.trsl.2014.01.005.

4. E. Tsolaki and E. Yannaki, “Stem cell-based regenerative opportunities for the liver: State of the art and beyond.,” World J. Gastroenterol., vol. 21, no. 43, pp. 12334–50, Nov. 2015, doi: 10.3748/wjg.v21.i43.12334.

5. F. Oldhafer, M. Bock, C. S. Falk, and F. W. R. Vondran, “Immunological aspects of liver cell transplantation.,” World J. Transplant., vol. 6, no. 1, pp. 42–53, Mar. 2016, doi: 10.5500/wjt.v6.i1.42.

6. P. Godoy et al., “Recent advances in 2D and 3D in vitro systems using primary hepatocytes, alternative hepatocyte sources and non-parenchymal liver cells and their use in investigating mechanisms of hepatotoxicity, cell signaling and ADME,” Arch. Toxicol., vol. 87, no. 8, pp. 1315–1530, Aug. 2013, doi: 10.1007/s00204-013-1078-5.

7. K. Takahashi and S. Yamanaka, “Induction of Pluripotent Stem Cells from Mouse Embryonic and Adult Fibroblast Cultures by Defined Factors,” Cell, vol. 126, no. 4, pp. 663–676, Aug. 2006, doi: 10.1016/J.CELL.2006.07.024.

8. K. Takahashi and S. Yamanaka, “Induced pluripotent stem cells in medicine and biology.,” Development,vol. 140, no. 12, pp. 2457–61, Jun. 2013, doi: 10.1242/dev.092551.

9. D. C. Hay et al., “Highly efficient differentiation of hESCs to functional hepatic endoderm requires ActivinA and Wnt3a signaling,” Proc. Natl. Acad. Sci. U. S. A., vol. 105, no. 34, p. 12301, 2008, doi: 10.1073/PNAS.0806522105.

10. J. Shan et al., “HHS Public Access,” vol. 9, no. 8, pp. 514–520, 2014, doi: 10.1038/nchembio.1270.Identification.

11. D. S. Varghese, T. T. Alawathugoda, and S. A. Ansari, “Fine Tuning of Hepatocyte Differentiation from Human Embryonic Stem Cells: Growth Factor vs. Small Molecule-Based Approaches.,” Stem Cells Int., vol. 2019, p. 5968236, 2019, doi: 10.1155/2019/5968236.

12. P. Matz, W. Wruck, B. Fauler, D. Herebian, T. Mielke, and J. Adjaye, “Footprint-free human fetal foreskin derived iPSCs: A tool for modeling hepatogenesis associated gene regulatory networks,” Sci. Rep., vol. 7, no. 1, p. 6294, Dec. 2017, doi: 10.1038/s41598-017-06546-9.

13. H. Rashidi et al., “3D human liver tissue from pluripotent stem cells displays stable phenotype in vitro and supports compromised liver function in vivo,” Arch. Toxicol., vol. 92, no. 10, pp. 3117–3129, Oct. 2018, doi: 10.1007/s00204-018-2280-2.

14. M. Kapałczyńska et al., “2D and 3D cell cultures - a comparison of different types of cancer cell cultures.,” Arch. Med. Sci., vol. 14, no. 4, pp. 910–919, Jun. 2018, doi: 10.5114/aoms.2016.63743.

15. G. A. Brent, “Mechanisms of thyroid hormone action.,” J. Clin. Invest., vol. 122, no. 9, pp. 3035–43, Sep. 2012, doi: 10.1172/JCI60047.

16. M. Bohndorf, A. Ncube, L.-S. Spitzhorn, J. Enczmann, W. Wruck, and J. Adjaye, “Derivation and characterization of integration-free iPSC line ISRM-UM51 derived from SIX2-positive renal cells isolated from urine of an African male expressing the CYP2D6 *4/*17 variant which confers intermediate drug metabolizing activity,” Stem Cell Res., vol. 25, 2017, doi: 10.1016/j.scr.2017.10.004.

17. J. Jozefczuk, A. Prigione, L. Chavez, and J. Adjaye, “Comparative Analysis of Human Embryonic Stem Cell and Induced Pluripotent Stem Cell-Derived Hepatocyte-Like Cells Reveals Current Drawbacks and Possible Strategies for Improved Differentiation,” Stem Cells Dev., vol. 20, no. 7, pp. 1259–1275, Jul. 2011, doi: 10.1089/scd.2010.0361.

18. N. Graffmann et al., “Modeling Nonalcoholic Fatty Liver Disease with Human Pluripotent Stem Cell-Derived Immature Hepatocyte-Like Cells Reveals Activation of PLIN2 and Confirms Regulatory Functions of Peroxisome Proliferator-Activated Receptor Alpha,” Stem Cells Dev., vol. 25, no. 15, 2016, doi: 10.1089/scd.2015.0383.

19. R. C. Gentleman et al., “Bioconductor: open software development for computational biology and bioinformatics.,” Genome Biol., vol. 5, no. 10, 2004, doi: 10.1186/gb-2004-5-10-r80.

20. B. S. Carvalho and R. A. Irizarry, “A framework for oligonucleotide microarray preprocessing,” Bioinformatics, vol. 26, no. 19, pp. 2363–2367, 2010, doi: 10.1093/bioinformatics/btq431.

21. H. Chen and P. C. Boutros, “VennDiagram: A package for the generation of highly-customizable Venn and Euler diagrams in R,” BMC Bioinformatics, vol. 12, no. 1, p. 35, 2011, doi: 10.1186/1471-2105-12-35.

22. M. Kanehisa, M. Furumichi, M. Tanabe, Y. Sato, and K. Morishima, “KEGG: New perspectives on genomes, pathways, diseases and drugs,” Nucleic Acids Res., vol. 45, no. D1, pp. D353–D361, 2017, doi: 10.1093/nar/gkw1092.

23. W. Luo and C. Brouwer, “Pathview: An R/Bioconductor package for pathway-based data integration and visualization,” Bioinformatics, vol. 29, no. 14, pp. 1830–1831, 2013, doi: 10.1093/bioinformatics/btt285.

24. S. Falcon and R. Gentleman, “Using GOstats to test gene lists for GO term association,” Bioinformatics, vol. 23, no. 2, pp. 257–258, 2007, doi: 10.1093/bioinformatics/btl567.

25. L. Burri, G. H. Thoresen, and R. K. Berge, “The role of PPAR activation in liver and muscle,” PPAR Res., vol. 2010, 2010, doi: 10.1155/2010/542359.

26. A. Flores-Morales et al., “Patterns of liver gene expression governed by TRβ,” Mol. Endocrinol., vol. 16, no. 6, pp. 1257–1268, 2002, doi: 10.1210/me.16.6.1257.

27. C. Lu and S.-Y. Cheng, “Thyroid hormone receptors regulate adipogenesis and carcinogenesis via crosstalk signaling with peroxisome proliferator-activated receptors,” J. Mol. Endocinology, vol. 44, no. 3, pp. 143–154, 2010, [Online]. Available: https://jme.bioscientifica.com/view/journals/jme/44/3/143.xml.

28. G. Y. Rychkov and G. J. Barritt, “Expression and Function of TRP Channels in Liver Cells BT - Transient Receptor Potential Channels,” M. S. Islam, Ed. Dordrecht: Springer Netherlands, 2011, pp. 667–686.

29. T. A. Potapova, V. N. Babenko, V. F. Kobzev, A. G. Romashchenko, V. N. Maksimov, and M. I. Voevoda, “Associations of Cold Receptor TRPM8 Gene Single Nucleotide Polymorphism with Blood Lipids and Anthropometric Parameters in Russian Population,” Bull. Exp. Biol. Med., vol. 157, no. 6, pp. 757–761, 2014, doi: 10.1007/s10517-014-2660-4.

30. M. Fortier et al., “Hepatospecific ablation of p38α MAPK governs liver regeneration through modulation of inflammatory response to CCl4-induced acute injury,” Sci. Rep., vol. 9, no. 1, pp. 1–12, 2019, doi: 10.1038/s41598-019-51175-z.

31. A. M. Tormos et al., “P38α regulates actin cytoskeleton and cytokinesis in hepatocytes during development and aging,” PLoS One, vol. 12, no. 2, pp. 1–22, 2017, doi: 10.1371/journal.pone.0171738.

32. M. Gordillo, T. Evans, and V. Gouon-Evans, “Orchestrating liver development,” Dev., vol. 142, no. 12, pp. 2094–2108, 2015, doi: 10.1242/dev.114215.

33. I. A. Rowe, “Lessons from Epidemiology: The Burden of Liver Disease,” Dig. Dis., vol. 35, no. 4, pp. 304–309, 2017, doi: 10.1159/000456580.

34. R. Siller, S. Greenhough, E. Naumovska, and G. J. Sullivan, “Small-Molecule-Driven Hepatocyte Differentiation of Human Pluripotent Stem Cells,” Stem Cell Reports, vol. 4, no. 5, pp. 939–952, May 2015, doi: 10.1016/j.stemcr.2015.04.001.

35. Y. Song, X. Yao, and H. Ying, “Thyroid hormone action in metabolic regulation.,” Protein Cell, vol. 2, no. 5, pp. 358–68, May 2011, doi: 10.1007/s13238-011-1046-x.

36. R. Malik and H. Hodgson, “The relationship between the thyroid gland and the liver,” QJM, vol. 95, no. 9, pp. 559–569, Sep. 2002, doi: 10.1093/qjmed/95.9.559.

37. Y.-H. Huang, M.-M. Tsai, and K.-H. Lin, “Thyroid Hormone Dependent Regulation of Target Genes and Their Physiological Significance,” 2008. Accessed: Apr. 26, 2019. [Online]. Available: http://cgmj.cgu.edu.tw/3104/310401.pdf.

38. R. Conti, C. Ceccarini, and M. F. Tecce, “Thyroid hormone effect on alpha-fetoprotein and albumin coordinate expression by a human hepatoma cell line.,” Biochim. Biophys. Acta, vol. 1008, no. 3, pp. 315–21, Aug. 1989, Accessed: Aug. 31, 2018. [Online]. Available: http://www.ncbi.nlm.nih.gov/pubmed/2474323.

39. Ö. Tarım, “Thyroid hormones and growth in health and disease.,” J. Clin. Res. Pediatr. Endocrinol., vol. 3, no. 2, pp. 51–5, 2011, doi: 10.4274/jcrpe.v3i2.11.

40. A. Cvoro et al., “A thyroid hormone receptor/KLF9 axis in human hepatocytes and pluripotent stem cells.,” Stem Cells, vol. 33, no. 2, pp. 416–28, Feb. 2015, doi: 10.1002/stem.1875.

