## Supplementary figures for "Investigation of Thyroid Hormone Associated Gene-Regulatory Networks during Hepatogenesis using an Induced Pluripotent Stem Cell based Model"

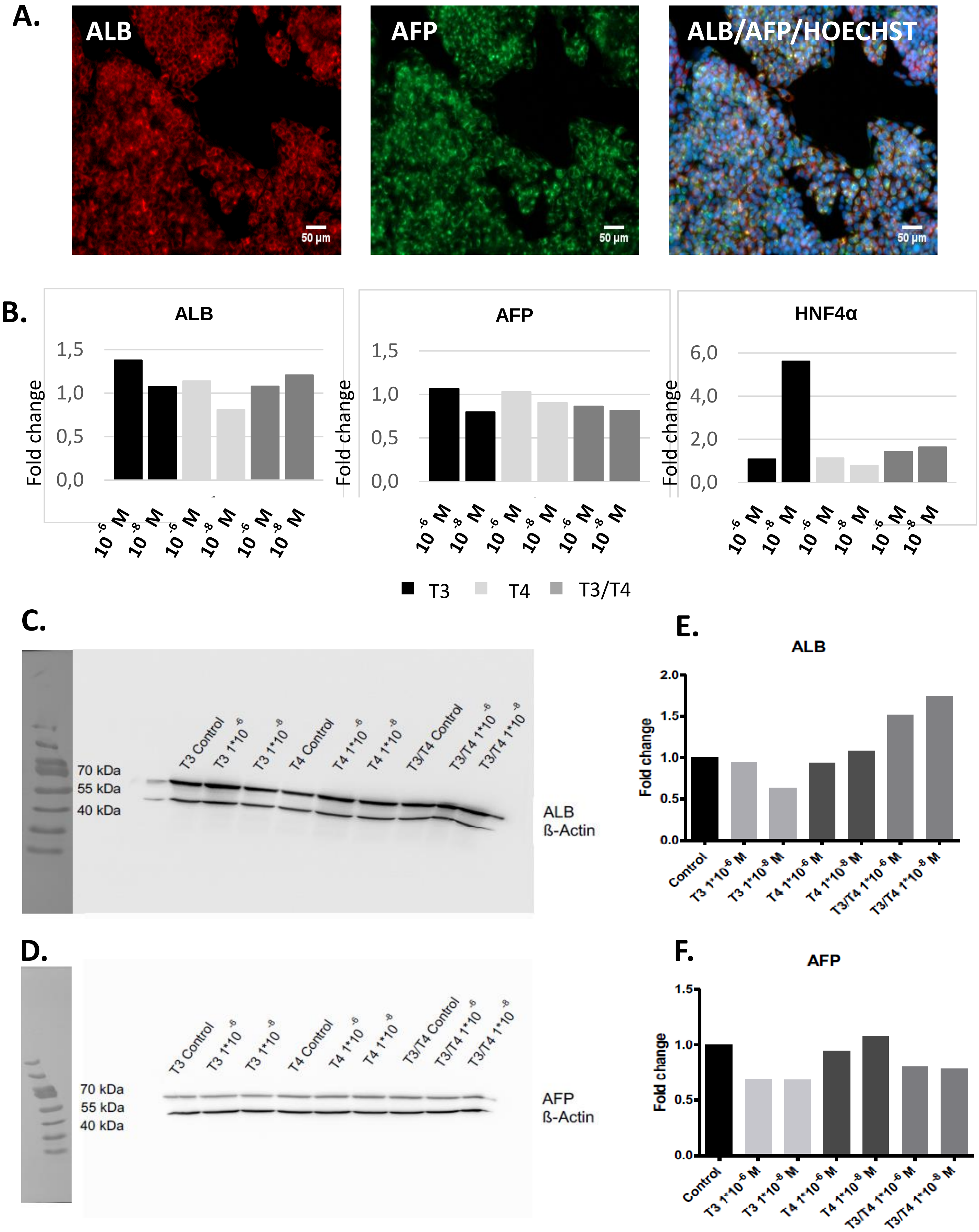
A.
ALB
AFP
ALB/AFP/HOECHST
50 µm
B.
ALB
AFP
HNF4α
Fold change
1,5
1,0
0,5
0,0
6,0
4,0
2,0
10 -6 M
10 -8 M
T3
T4
T3/T4
C.
70 kDa
55 kDa
40 kDa
T3 Control
T3 1*10 -6
T3 1*10 -8
T4 Control
T4 1*10 -6
T4 1*10 -8
T3/T4 Control
T3/T4 1*10 -6
T3/T4 1*10 -8
ALB
ß-Actin
D.
AFP
ß-Actin
E.
F.
2.0
1.5
1.0
0.5
0.0
Control
T3 1*10-6 M
T3 1*10-8 M
T4 1*10-6 M
T4 1*10-8 M
T3/T4 1*10-6 M
T3/T4 1*10-8 M

**Supplementary Figure S1.** Dosis determination of T3 and T4 after 7 days treatment of HepG2 cells. (A) Immunofluorescence staining images of the different conditions after 7 days treatment of the HepG2 cells. The cells were stained for ALB, AFP and the nuclei staining Hoechst. Untreated cells were compared to those treated with either T3, T4 or the combination T3/T4 at concentrations $10^{-6}$ M and $10^{-8}$ M. There were no visible differences in the staining images of the different treatment conditions nor concentrations. (B) qRT-PCR evaluation comparing the 2–ddCT values from the amplification. The mean CT values were normalised to the housekeeping gene RPS16 and the fold change evaluated relative to the untreated control cells. A slight increase in ALB expression was seen under T3 and the combination T3/T4 treatment in both concentrations $10^{-6}$ M and $10^{-8}$ M. An HNF4$\alpha$ high increase was observed after T3 treatment. (C, D) Western blot for ALB (C) and AFP (D) with the indicated concentrations of T3 and/or T4. (E, F) The band pixel intensities were normalised to ß-Actin and fold change evaluated relative to the controls. ALB expression increased upon T3/T4 treatment, whereas an AFP expression decrease was observed after T3 and T3T4 treatment in both concentrations.

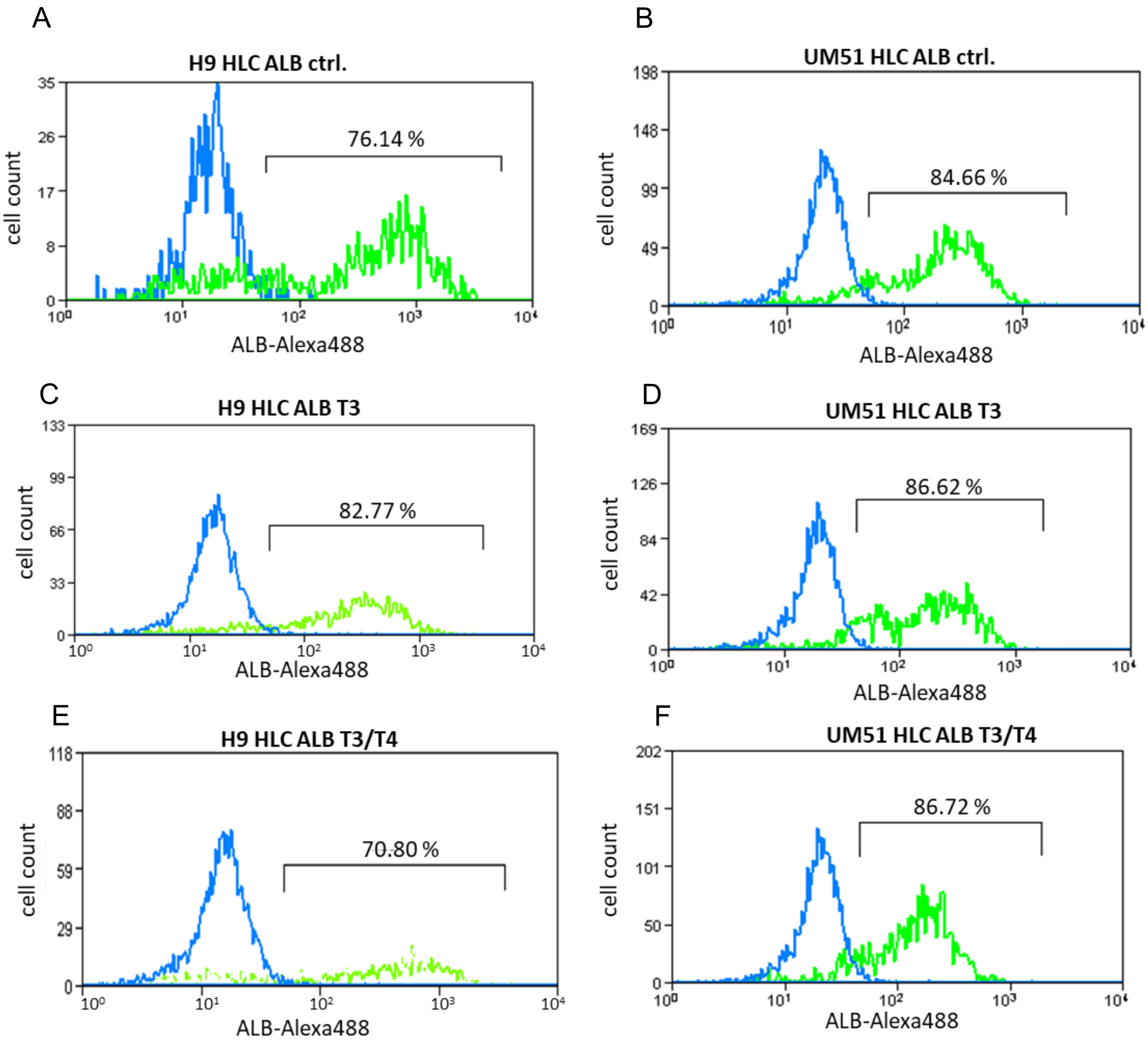


**Supplementary Figure S2:** Representative flow cytometry analysis of ALB expression in HLCs after cultivation in standard conditions (A,B), with T3 alone (C,D) or T3/T4 (E,F). Blue curve: secondary antibody control, green curve: ALB staining.

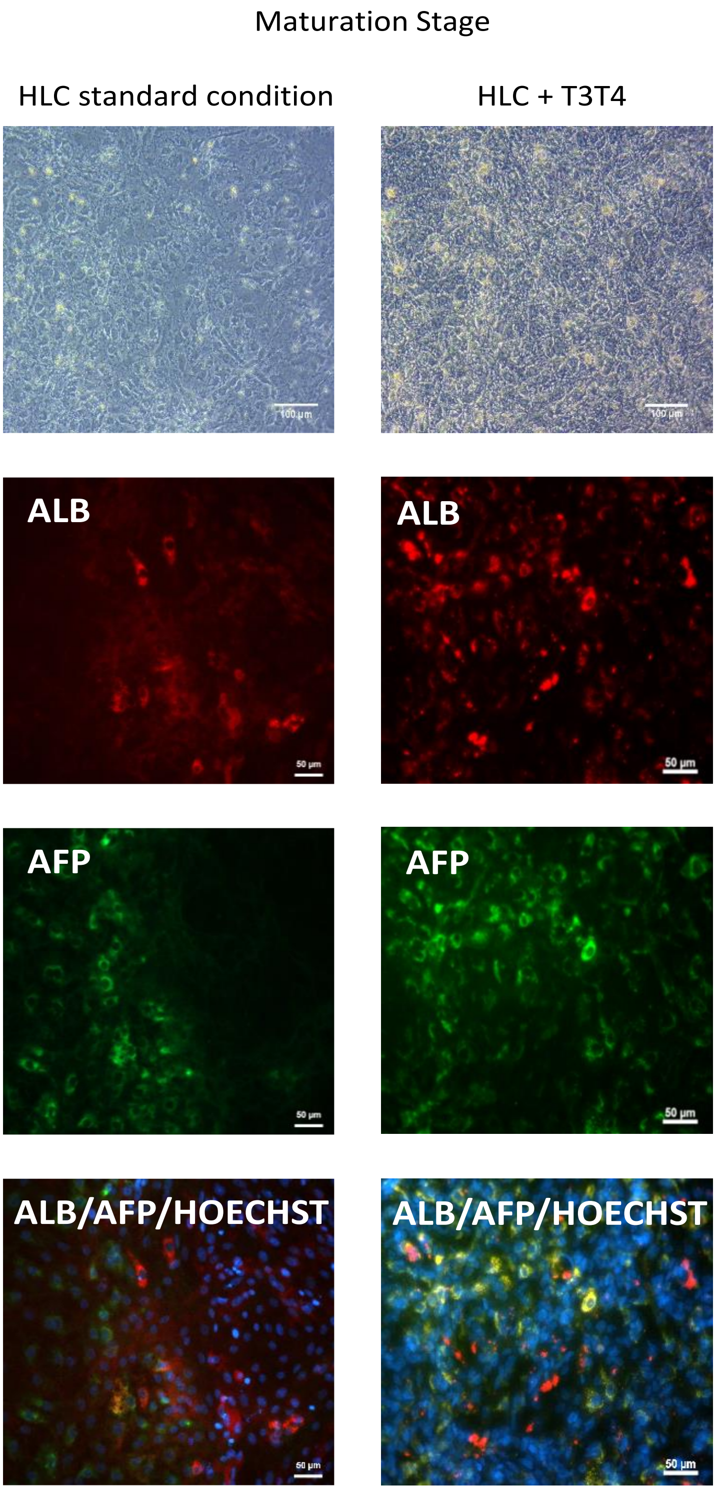


**Supplementary Figure S3.** Immunostaining images of HLCs generated under standard condition compared to those generated under T3T4 treatment, showing ALB and AFP staining. (Bright-field images Scale bars = 100 μm, immunostaining images scale bars = 50 μm).

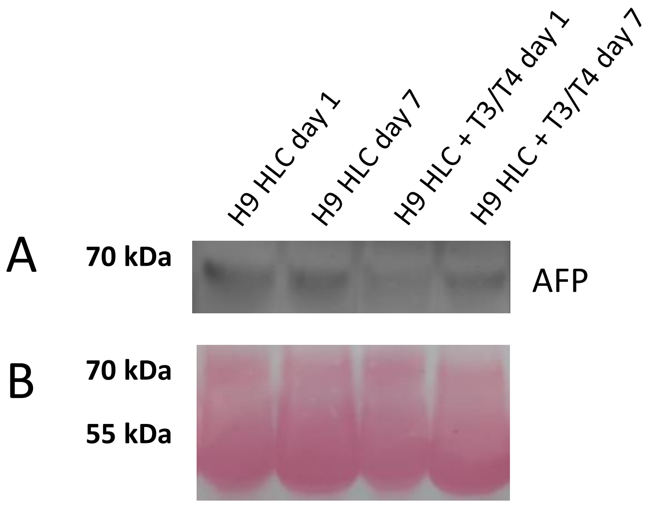


**Supplementary Figure S4:** Western blot was performed with concentrated supernatants from ESC derived HLCs. (A) AFP could be detected in the supernatant. (B) Ponceau S staining.

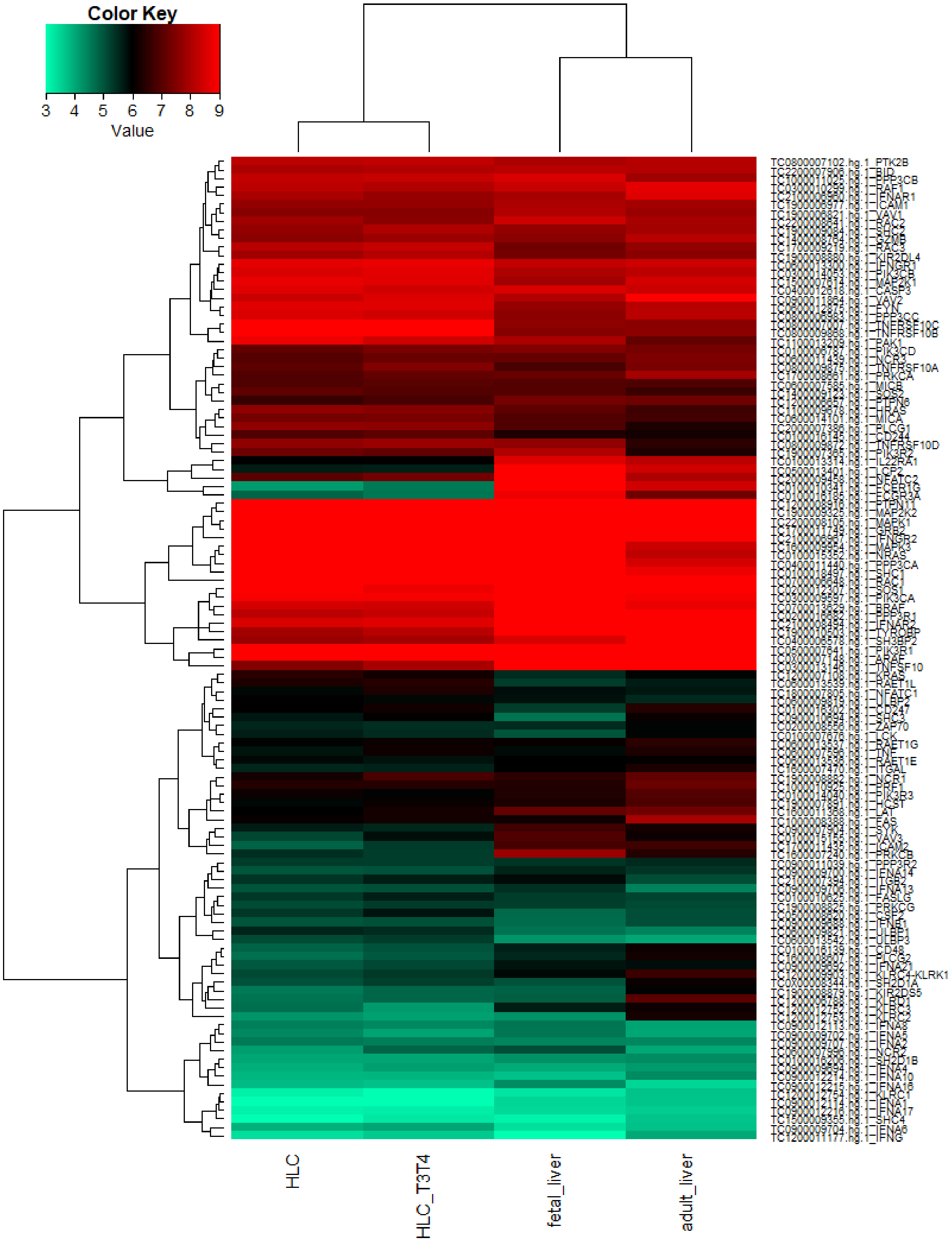


**Supplementary Figure S5:** Genes from the KEGG pathway "Natural killer cell mediated cytotoxicity" (hsa04650) are not affected by the T3/T4 treatment.

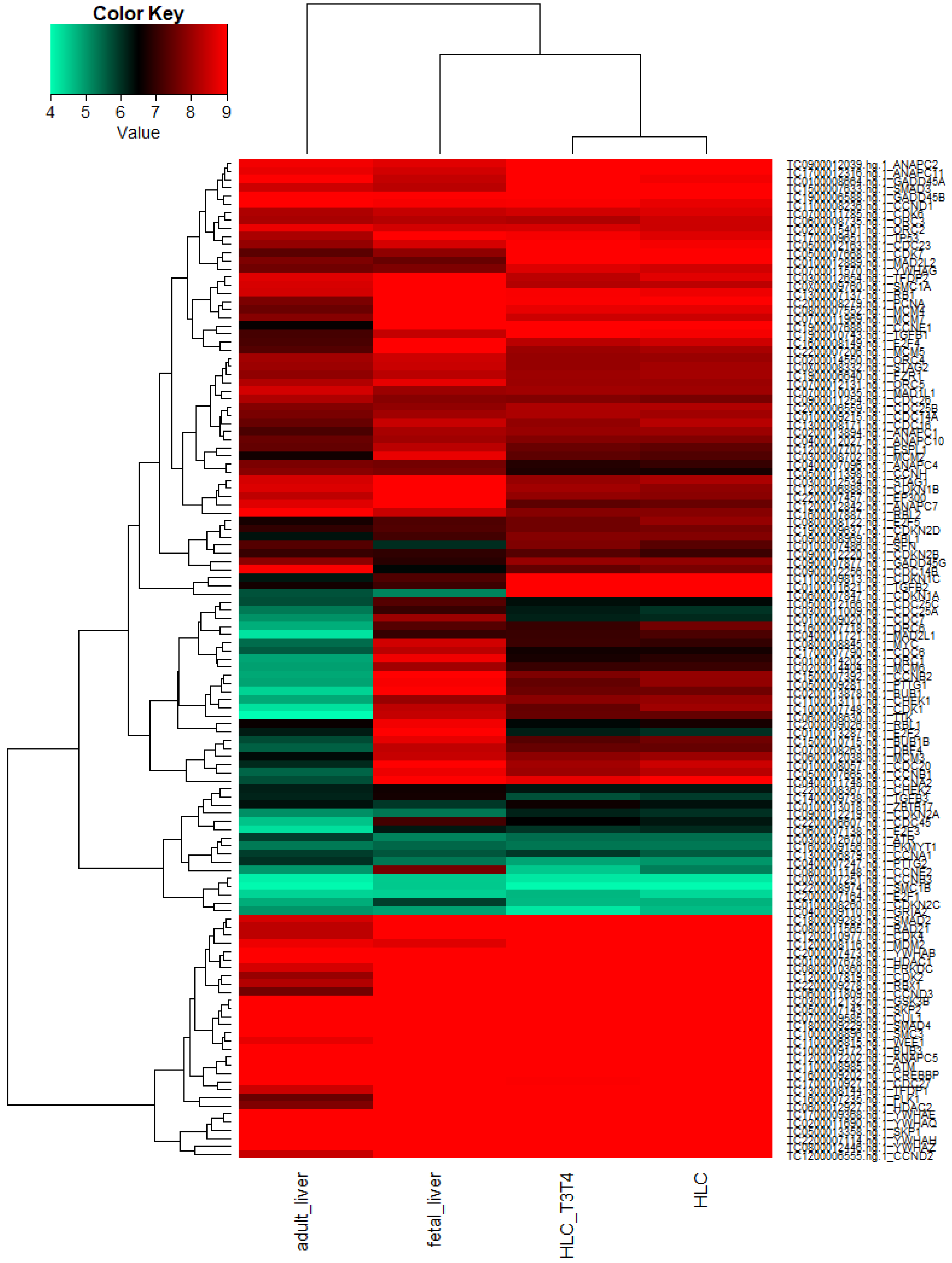


**Supplementary Figure S6:** Genes from the KEGG pathway "Cell cycle" (hsa04110) are not or only marginally affected by the T3/T4 treatment.
