## Supplementary Table S2 for "Investigation of Thyroid Hormone Associated Gene-Regulatory Networks during Hepatogenesis using an Induced Pluripotent Stem Cell based Model"

**Supplementary Table S2:** List of specific primer sequences.

| **Primer** |  | **Sequence (5‘→3‘)** |
| --- | --- | --- |
| AFP | Forward | AGCAGCTTGGTGGTGGATGA |
|  | Reverse | CCTGAGCTTGGCACAGATCCT |
| ALBUMIN | Forward | AGCTGTTATGGATGATTTCGCAG |
|  | Reverse | CCTCGGCAAAGCAGGTCTC |
| CYP3A4 | Forward | GTGACTTTGCCCATTGTTTAGAAAG |
|  | Reverse | CAGGCGTGAGCCACTGTG |
| RPS16 | Forward | GCTATCCGTCAGTCCATCTCCAA |
|  | Reverse | CCTTCTTGGAAGCCTCATCCAC |
| A1AT | Forward | AGCAGCTTGGTGGTGGATGA |
|  | Reverse | CCTGAGCTTGGCACAGATCCT |
| TTR | Forward | CTGCCTTGCTGGACTGGTAT |
|  | Reverse | CAGCAGCCTTTCTGAACACA |
| CYP3A7 | Forward | GATTCTGTACGTGCATTGTGCTC |
|  | Reverse | ATTTGGTCATCTCCTCTATATTACCAAGT |
